# Entorhinal cortex epigenome-wide association study highlights four novel loci showing differential methylation in Alzheimer’s disease

**DOI:** 10.1101/2021.07.02.450878

**Authors:** Yasmine Sommerer, Valerija Dobricic, Marcel Schilling, Olena Ohlei, Sanaz Sedghpour Sabet, Tanja Wesse, Janina Fuß, Sören Franzenburg, Andre Franke, Laura Parkkinen, Christina M. Lill, Lars Bertram

## Abstract

**Background:** Studies on DNA methylation (DNAm) in Alzheimer’s disease (AD) have recently highlighted several genomic loci showing association with disease onset and progression.

**Methods:** Here, we conducted an epigenome-wide association study (EWAS) using DNAm profiles in entorhinal cortex (EC) from 149 AD patients and control brains and combined these with two previously published EC datasets by meta-analysis (total n=337).

**Results:** We identified 12 cytosine-phosphate-guanine (CpG) sites showing epigenome-wide significant association with either case-control status or Braak’s tau-staging. Four of these CpGs, located in proximity to *CNFN/LIPE, TENT5A, PALD1/PRF1*, and *DIRAS1*, represent novel findings. Integrating DNAm levels with RNA sequencing-based mRNA expression data generated in the same individuals showed significant DNAm-mRNA correlations for 6 of the 12 significant CpGs. Lastly, by calculating rates of epigenetic age acceleration using two recently proposed “epigenetic clock” estimators we found a significant association with accelerated epigenetic aging in AD patients vs. controls.

**Conclusion:** In summary, our study represents the hitherto most comprehensive EWAS in AD using EC and highlights several novel differentially methylated loci with potential effects on gene expression.

## Introduction

Alzheimer’s disease (AD) is a progressive, neurodegenerative disease that accounts for 50-60% of all dementia cases (1). The number of AD cases is increasing and it was recently estimated that nearly 44 million individuals lived with dementia in 2016 world-wide (2). On a neuropathological level, the hallmarks of AD are accumulations of amyloid-beta (Aβ) plaques and neurofibrillary tangles (NFTs) consisting of hyperphosphorylated tau protein. There is growing evidence that the first neuropathological changes already occur two decades or more prior to the onset of clinical symptoms (3). In early AD, NFTs are regularly observed without the formation of Aβ, with neuropathological changes typically starting in the transentorhinal followed by the entorhinal cortex (EC) before spreading across most cortical brain regions while the disease progresses (4). Owing to this spatio-temporal course, the EC represents an interesting and informative brain region to study in molecular AD research, including studies aimed at the epigenome or transcriptome.

There is accumulating evidence that epigenetic factors (in addition to genetic factors) may contribute to the onset and progression of AD (5–7). One of the most widely studied epigenetic marks is DNA methylation (DNAm) owing to the relative technical ease to generate these data on a(n) (epi)genome-wide scale. Since 2014, this has led to a number of epigenome-wide association studies (EWAS) assessing DNAm profiles in various AD-related phenotypes (7–16), culminating in a very recent meta-analysis on differential DNAm across various brain datasets (17). Taken together, these studies identified several genomic loci (e.g., *ANK1, RPL13, SPG7*, and *MCF2L*) showing consistent changes in DNAm patterns associated with AD related phenotypes across several brain regions (e.g., EC as well as temporal and prefrontal cortex).

In this study, we generated DNAm (using the MethylationEPIC microarray) and mRNA (using RNA sequencing) expression profiles in the same EC slices from 65 AD cases and 84 control brains. These data were used to conduct a DNAm-based EWAS using both case-control status and Braak’s tau-staging (henceforth termed “Braak staging”) as predictors. For the EWAS part, we combined our DNAm data with data from two previously published EC studies (both generated using the 450K Methylation microarray) (11,17) increasing our total sample size to n=337. Significantly differentially methylated sites were then correlated with corresponding mRNA levels to probe for potential effects of DNAm on gene expression.

## Methods and Materials

### Human samples

Snap-frozen, post-mortem human brain tissue from EC slices (Brodmann area BA28) from 91 AD patients and 92 elderly control individuals was obtained from the Oxford Brain Bank. The AD patients and healthy controls were part of the longitudinal, prospective Oxford Project to Investigate Memory and Aging (OPTIMA) using protocols which have been described in detail elsewhere (18). All subjects underwent a detailed clinical history, physical examination, assessment of cognitive function (Cambridge Examination of Mental Disorders of the Elderly (CAMDEX) (19) with the Cambridge Cognitive Examination (CAMCOG) and Mini-Mental State Examination (MMSE) biannually. The pathological diagnosis of AD was made using the Consortium to Establish a Registry for Alzheimer’s disease (CERAD)/National Institutes of Health (NIH) criteria and Braak staging (20–22). All included patients were of white European descent by self-report. The Ethics Committees of Oxford University and University of Lübeck approved the use of the human tissues for our study and all participants gave informed consent. Details regarding the DNA and RNA extraction, as well as the procedures for DNAm profiling using the “Infinium MethylationEPIC” array (EPIC; Illumina, Inc.), RNA sequencing, and quality control (QC) can be found in the supplementary methods. The EPIC array is the successor to the widely used “Infinium HumanMethylation450” kit; the high precision and reproducibility of the DNAm data generated with the EPIC array has been validated in a large number of independent reports (23–25). A detailed sample description can be found in Table 1.

**Table 1:** Demographic data for the entorhinal cortex Oxford dataset by case-control status (n=149) and Braak staging (n=142) after QC.

| AD Status / Braak Stage | Sample size<br>(% AD) | % Females | Age / years |
| --- | --- | --- | --- |
| Control | 65 | 40 | $80 \pm 13$ |
| Case | 84 | 51 | $82 \pm 8$ |
| Braak stage 0 | 3 (0) | 33 | $66 \pm 4$ |
| Braak stage I | 8 (0) | 25 | $74 \pm 11$ |
| Braak stage I/II | 27 (0) | 37 | $85 \pm 8$ |
| Braak stage II | 10 (0) | 40 | $81 \pm 11$ |
| Braak stage III | 2 (0) | 100 | $97 \pm 5$ |
| Braak stage III/IV | 8 (13) | 50 | $87 \pm 3$ |
| Braak stage IV | 6 (83) | 17 | $88 \pm 4$ |
| Braak stage V | 24 (100) | 29 | $86 \pm 6$ |
| Braak stage V/VI | 24 (100) | 67 | $81 \pm 9$ |
| Braak stage VI | 30 (100) | 60 | $88 \pm 7$ |

### Epigenome-wide association study (EWAS) analyses to identify differentially methylated probes (DMPs) and differentially methylated regions (DMRs)

Statistical analyses to identify differentially methylated probes (DMPs) were performed based on linear regression models using the *lm* function in R using case-control status (as dichotomous variable) or Braak stage (as continuous variable) as predictor in the EWAS, respectively:

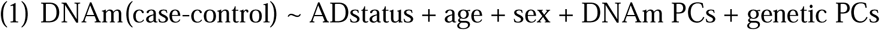

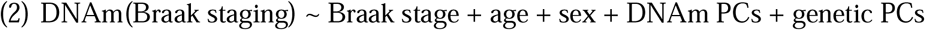

To account for differences in the DNAm profiles due to technical (e.g. laboratory batch, array) and other (e.g. cell-type composition of samples, genetic ancestry) factors we implemented an elaborate protocol of batch effect correction (see Supplement for full details).

Differentially methylated regions (DMRs, i.e. combinations of consecutive DMPs) were assessed with the comb-p tool (26) with the maximal gap within a region set to 500 base pairs and the seed *p*-value set to 1.00E-03. Only regions including at least three cytosine-phosphate-guanine (CpG) sites were considered. Significance of the DMR results was determined using the Sidak method (as implemented in comb-p).

Annotation of CpGs to specific gene regions was based on the Illumina manifest (v1.0 B5) for the EPIC array and the GREAT annotation tool (27).

### Meta-analysis of epigenome-wide association study (EWAS) results

To increase power of our EWAS, we combined our EPIC array-based results with those from two publicly available AD EC datasets (GEO accession numbers GSE59685 with 58 AD cases and 21 controls [“London-1”]; and GSE105109 with 68 AD cases and 28 controls [“London-2”]). The descriptions of these datasets can be found in the primary publications (7,11). Here, we downloaded the processed DNAm values and repeated EWAS analyses for Braak stage and AD case-control status using the same linear regression models as described above. These regression models included 13 DNAm PCs for the Braak stage analysis of GSE59685, and 15 DNAm PCs for the remaining analyses. The meta-analysis was conducted with a fixed-effect inverse-variance approach using the function *metagen* in the R package “meta” (29).

### Alzheimer’s disease poly-epigenetic scores

To assess the correspondence of our novel DNAm data to previous EWAS on the topic, poly-epigenetic scores (PES) for each individual were calculated based on the test statistics from two publicly available AD EC datasets (GEO accession numbers GSE59685; GSE105109). To this end, we combined uncorrelated CpGs into one aggregated DNAm variable and tested these as predictors in regression models analogous to the primary EWAS. For more details see Supplementary Material.

### Epigenetic age estimation

Two epigenetic age predictors were used in our analyses: 1. the “Horvath multi-tissue predictor” (HMTP) (28) and 2. the “cortex clock” (CorCl) (30). Since most other popular epigenetic clocks (Hannum (31), PhenoAge (32), GrimAge (33)) were calibrated for blood tissues, we did not include analyses of these age estimators in this study. For more details regarding the calculation of the epigenetic age estimates see Supplementary Material.

### DNAm-mRNA correlation analyses

The normalized RNA-seq data for the selected CpG candidate genes (see above) were correlated to their corresponding DNAm signal, using the Spearman method in R’s *cor*.*test* function. The resulting *p*-values were corrected for multiple testing using the Benjamini-Hochberg procedure (as implemented in R’s *p*.*adjust* function). For more details, see Supplementary Material.

## Results

### EWAS of case-control status and Braak staging highlights five DMRs

In the EWAS analyses of all 665,796 CpG-probes that passed QC on the EPIC array, none of the CpGs reached the experiment-wide Bonferroni-corrected significance threshold α = 7.51E-08, neither in the analyses of AD case-control status (Supplementary Figure 2) nor Braak stage (Supplementary Figure 3). However, we note that several CpGs reached at least suggestive evidence of association with either phenotype (α = 1.00E-05; Supplementary Tables 1 and 2). Among these, cg25191519 showed a particularly strong association signal (*p* = 8.90E-06). This CpG is annotated to the genes *SPG7* and *RPL13*, which were already reported in previous AD DNAm studies (8,15–17). Next, we used the DMP test statistics to assess the presence of DMRs, i.e., consecutive runs of differentially methylated probes which are aggregated into “regions”. After adjustment for multiple testing, five DMRs (near genes *PRKCZ, CYFIP1, ACOT7, COL4A1, IBA57*, and *C1orf69*) showed significant association with AD case-control status (Table 2). In contrast, no DMRs were found when combining DMPs from the Braak stage EWAS using p-comb. Two of the six genes highlighted by the significant case-control DMRs were previously described in the context of AD DNAm profiling studies. First, a DMR near *CYFIP1* was reported by Li et al. to show association with Braak stage (15). Second, the gene *IBA57* is annotated to CpG-probe cg12461930, which also showed association with Braak stage in the cross-cortex meta-analysis by Smith et al. (17).

**Table 2:** Results of EWAS using AD case-control status in Oxford sample. AD case-control DMRs identified using p-comb software. CpGs: number of CpGs in each DMR; *p*_adj_: *p*-value after adjustment for multiple testing based on the Sidak method. † Evidence for implication of same or largely overlapping locus from studies using DNAm assessments in AD-related phenotypes. More (*p* < 1.00E-05) results from this analysis can be found in Supplementary Table 1 and 2.

| Position | Gene | CpGs | $p$-value | $p_{\text{adj}}$ | Previous studies (ref.)<sup>†</sup> |
| --- | --- | --- | --- | --- | --- |
| chr1:2004968-2005180 | *PRKCZ* | 3 | 4.50E-11 | 1.41E-07 |  |
| chr15:22921227-22921426 | *CYFIP1* | 3 | 4.75E-10 | 1.59E-06 | DMR in *CYFIP1* (15) |
| chr1:6445901-6445975 | *ACOT7* | 3 | 3.80E-10 | 3.42E-06 |  |
| chr13:110918122-110918331 | *COL4A1* | 3 | 1.82E-09 | 5.80E-06 |  |
| chr1:228362233-228362309 | *IBA57 / C1orf69* | 3 | 6.03E-08 | 5.30E-04 | *IBA57* previously described (17) |

To further evaluate the degree of correspondence of our novel EWAS data with those from the recent EWAS meta-analysis by Smith et al. (17), we repeated the Braak stage meta-analysis from two publicly available EC DNAm datasets used in previous AD EWAS (“London-1” and “London-2”, with GEO accession numbers GSE59685 and GSE105109, respectively) (7,11), which were also included in the Smith et al. study (17), and used these resulting test statistics with varying *p*-value thresholds to calculate the PES. We then tested the PES for association with Braak stage in our dataset using linear models equivalent to the primary EWAS. These analyses revealed that the PES was, indeed, significantly associated with Braak stage in our dataset (7.83E-06 ≤ *p* ≤ 1.88E-01, Supplementary Table 3) and explained up to 12% of the phenotypic variance in our dataset. Overall, these results suggest that our dataset is equivalent in terms of data quality when compared to those previously published in the field.

### EWAS meta-analysis highlights 12 DMPs showing experiment-wide significant association with AD

To increase power, we combined the EWAS results from our samples with those from the two publicly available EC DNAm datasets London-1 and London-2. Of note, the DNAm data of these prior studies were generated on the predecessor 450K DNAm array which has a substantially lower resolution leading to a smaller number of meta-analysed CpGs. The meta-analysis across all three datasets comprised 320 samples for the AD case-control analysis, and 337 samples for the Braak stage analysis. Overall, there were 304,996 overlapping CpGs available for this meta-analysis, resulting in an experiment-wide significance threshold of α = 1.64E-07 for this arm of our study. Using this threshold, five CpGs in the AD case-control meta-analysis (Figure 1, Table 3), and nine CpGs in the Braak stage meta-analysis (Figure 2, Table 4), reached experiment-wide significance. Importantly, four of these were not previously reported in the context of EWAS using AD Braak stage or case-control status as phenotypes and can be considered *bona fide* novel findings of our study. Two CpGs (cg03169557 [near the genes *RPL13* and *SPG7*] and cg05066959 [near the genes *NKX6-3, ANK1*, and *MIR486*]) were significantly associated with both AD case-control status and AD Braak stage and were already described in previous DNAm AD studies (8,15–17). While none of the analyses in the individual datasets analyses showed notable inflation in the test statistics (Supplementary Table 4), both meta-analyses displayed slightly increased inflation (λ_case-control_ = 1.16; λ_Braak_ = 1.24; Supplementary Figure 4), a relatively common observation in EWAS as already noted in Smith et al. (17).

**Figure 1:**
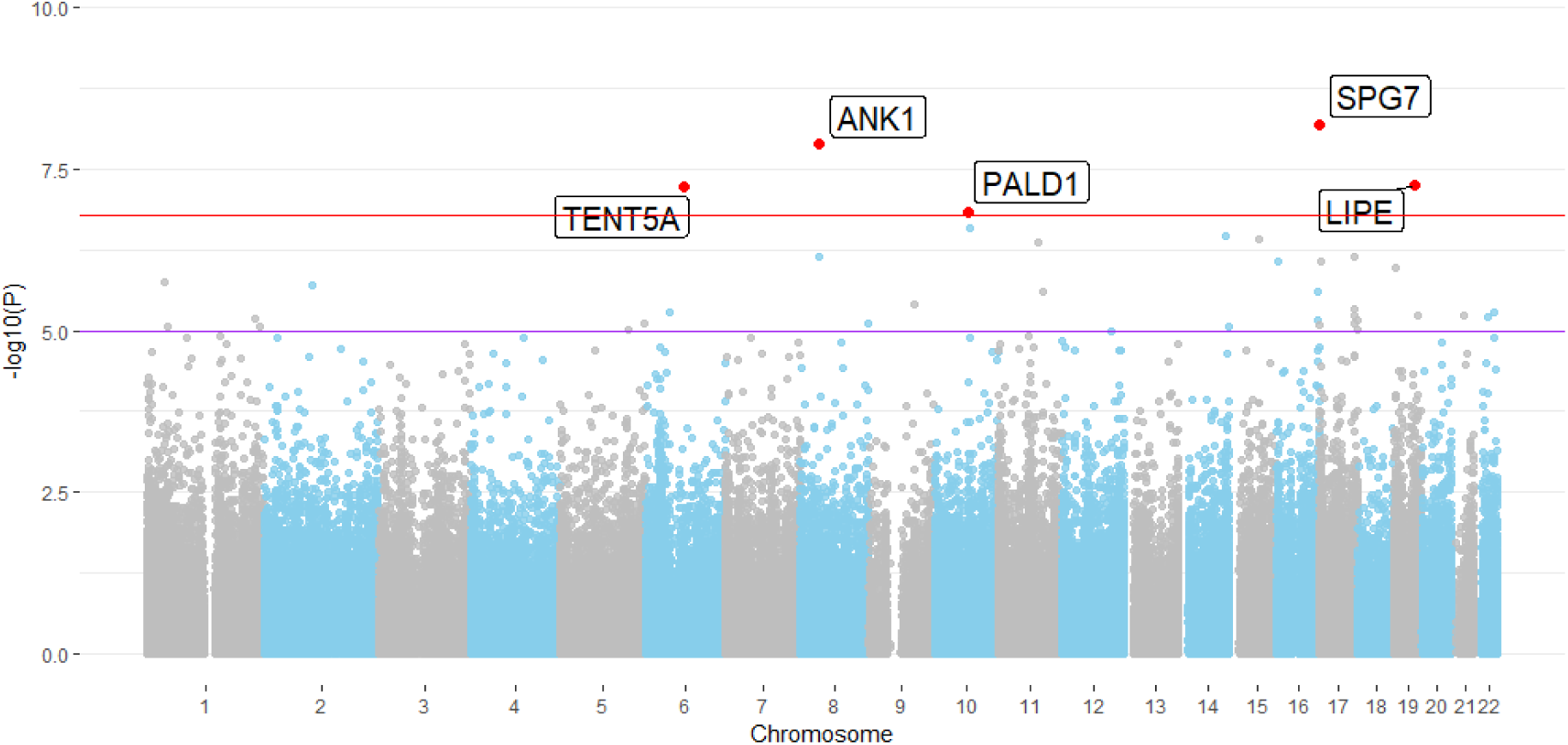
Manhattan plot for the EWAS meta-analysis using AD case-control status across three EC datasets. the red line indicates the experiment-wide significance threshold of 1.64E-07, whereas the purple line indicates the suggestive significance threshold of 1.00E-05. CpGs with experiment-wide significant association are marked in red and annotated with the gene name according to the Illumina manifest (v1.0 B5). CpGs around *LIPE, TENT5A, and PALD1* are novel.

**Table 3:** Results of EWAS meta-analysis using AD case-control status. Experiment-wide significant CpGs (*p* < 1.64E-07) in the meta-EWAS ascross three datasets (London-1, London-2, Oxford) using AD case-control status.† Evidence for implication of same or largely overlapping locus from studies using DNAm assessments in AD-related phenotypes. Annotation of CpGs to specific gene regions was based on the Illumina manifest (v1.0 B5) for the EPIC array and the GREAT annotation tool (27).

| CpG | Position | Genes | p-value | Effect ($\beta$) | Previous Studies (ref.)† |
| --- | --- | --- | --- | --- | --- |
| **cg03169557** | chr16:89598950 | *RPL13, SPG7* | 6.59E-09 | 0.0214 | (8,15–17) |
| **cg05066959** | chr8:41519308 | *NKX6-3, ANK1, MIR486* | 1.30E-08 | 0.0453 | (7,8,15–17) |
| **cg03073402** | chr19:42927676 | *CNFN, LIPE* | 5.59E-08 | -0.0086 | - |
| **cg22388948** | chr6:82460558 | *TENT5A* | 5.83E-08 | -0.0270 | - |
| **cg20648333** | chr10:72298745 | *PALD1, PRF1* | 1.44E-07 | -0.0183 | - |

**Figure 2:**
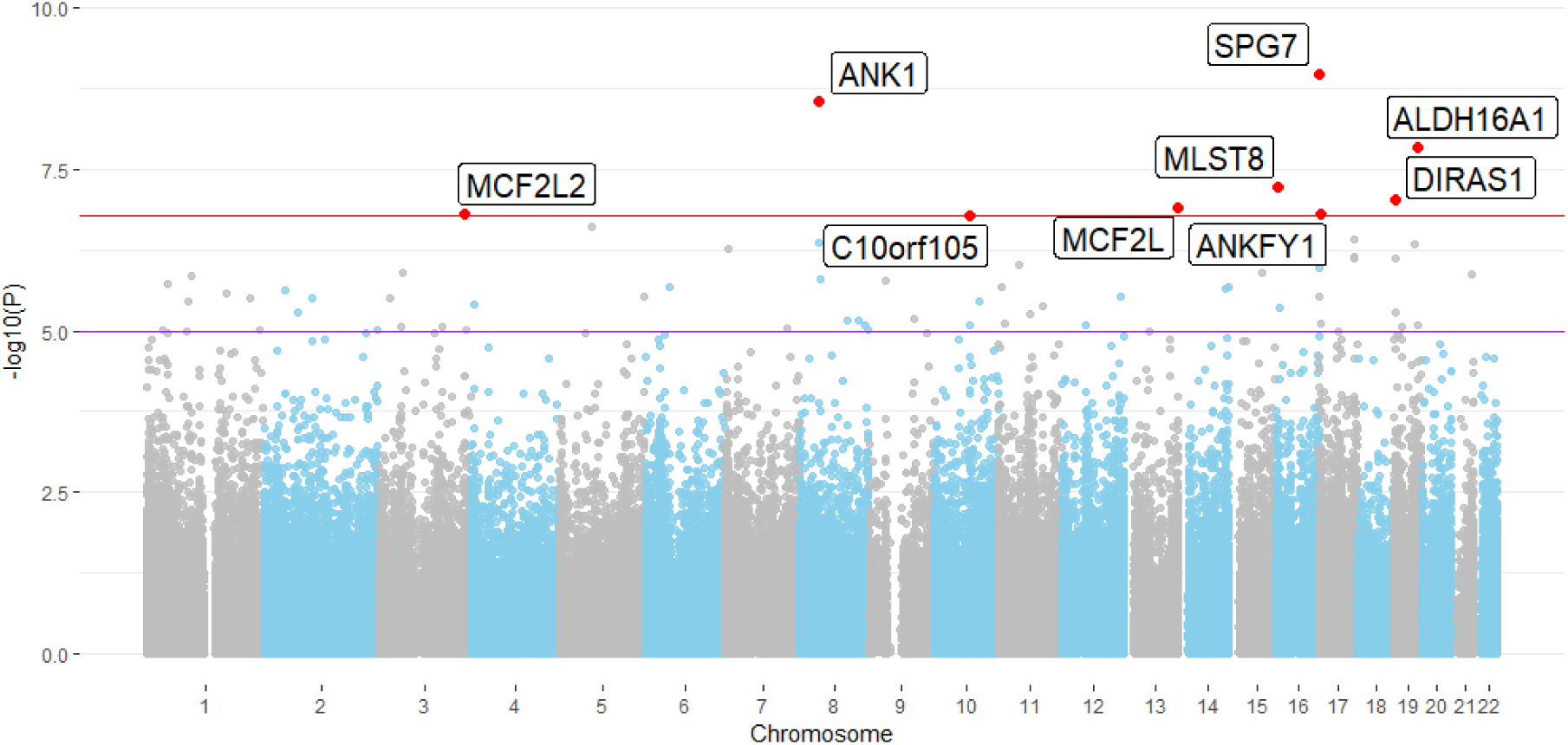
Manhattan plot for the EWAS meta-analysis using Braak staging across three EC datasets. the red line indicates the experiment-wide significance threshold of 1.64E-07, whereas the purple line indicates the suggestive significance threshold of 1.00E-05. CpGs with experiment-wide significant association are marked in red and annotated with the gene name according to the Illumina manifest (v1.0 B5). The CpG around *DIRAS1* is novel.

**Table 4:** Results of EWAS meta-analysis using Braak staging. Experiment-wide significant CpGs (*p* < 1.64E-07) in the meta-EWAS ascross three datasets (London-1, London-2, Oxford) using Braak stage. † Evidence for implication of same or largely overlapping locus from studies using DNAm assessments in AD-related phenotypes. Annotation of CpGs to specific gene regions was based on the Illumina manifest (v1.0 B5) for the EPIC array and the GREAT annotation tool (27)

| CpG | Position | Genes | p-value | Effect ($\beta$) | Previous Studies (ref.)† |
| --- | --- | --- | --- | --- | --- |
| **cg03169557** | chr16:89598950 | *RPL13, SPG7* | 1.03E-09 | 0.0042 | (8,15–17) |
| **cg05066959** | chr8:41519308 | *NKX6-3, ANK1, MIR486* | 2.76E-09 | 0.0092 | (7,8,15–17) |
| **cg20618448** | chr19:49962324 | *FLT3LG, ALDH16A1* | 1.42E-08 | 0.0043 | (17) |
| **cg05030077** | chr16:2255199 | *MLST8* | 5.89E-08 | -0.0026 | (11) |
| **cg05228284** | chr19:2720847 | *DIRAS1* | 9.25E-08 | -0.0023 | - |
| **cg05972352** | chr13:113663373 | *MCF2L, F2* | 1.20E-07 | 0.0058 | genes previously reported (8,10,15,17) |
| **cg14761246** | chr3:182968758 | *MCF2L2, B3GNT5* | 1.55E-07 | 0.0039 | (16) |
| **cg22090150** | chr17:4098227 | *ANKFY1, CYB5D2* | 1.57E-07 | 0.0048 | (13,16,17) |
| **cg07571519** | chr10:73472315 | *C10orf105, SLC29A3* | 1.62E-07 | 0.0050 | (16) |

The four newly associated CpG-probes cg03073402, cg22388948, cg20648333, and cg05228284 are located in or near the genes *CNFN*/*LIPE, TENT5A, PALD1/PRF1*, and *DIRAS1*, respectively. The implicated CpGs all displayed a reduction of DNAm associated with AD, i.e., a negative effect size estimate. Of these genes, *DIRAS1* shows the most pronounced expression in brain (GTEx V8) followed by *LIPE, TENT5A*, and *PALD1*. In contrast, *CNFN* and *PRF1* do not show any noteworthy expression in the brain tissues analysed in GTEx. Using a PubMed search (using “{gene name} AND alzheimer*} as search terms) revealed that no publication exists to date directly linking any of these genes to AD. Look-up of the CpG IDs on the EWAS Catalog (http://www.ewascatalog.org/) (34) revealed that cg03073402 (*CNFN/LIPE*) and cg20648333 (*PALD1/PRF1*) were previously reported to be associated with aging from birth to late adolescence in blood samples (35).

The GWAS catalog (https://www.ebi.ac.uk/gwas/ (36)) also revealed no noteworthy AD-related entries for *CNFN, LIPE, PALD1, PRF1*, or *DIRAS1*. In contrast, genetic variants in *TENT5A* (a.k.a. *FAM46A*) have been found associated by GWAS with a number of traits (https://www.ebi.ac.uk/gwas/genes/TENT5A), some of them with direct relevance for AD, e.g. “Alzheimer’s disease, posterior cortical atrophy”, “Alzheimer’s disease, cognitive decline measurement”, “PHF-tau measurement”, “neurofibrillary tangles measurement”, and “temporal pole volume measurement”, underscoring the potential mechanistic involvement of this gene in AD pathogenesis. The EWAS signal for *TENT5A* was elicited by CpG-probe cg22388948 (located intronically) with a *p*-value of 5.83E-08 in the AD case-control meta-analysis. Suggestive evidence for association with AD case-control status with the same effect direction could also be observed in the individual London datasets (*p*_L1_ = 6.73E-05, effect_L1_ = -0.0375, *p*_L2_ = 1.72E-02, effect_L2_ = -0.0174), emphasizing the robustness of the signals across independent datasets. While the remaining eight epigenome-wide significant CpGs of our meta-analyses were not featured as “top results” in the Smith et al. paper, they were highlighted in other previous AD EWAS (see Table 3 and 4, Supplementary Table 5).

Interestingly, three of the eight CpG probes that were previously reported to show experiment-wide significant association with Braak stage in EC (17) and were also present in our analysis did not show any evidence of association with Braak stage (*p* < 0.05) in the Oxford dataset (Supplementary Table 6). This relates to CpG probes cg04523589 (annotated to the gene *CAMP*), cg06653632 (annotated to *SLC15A4* and *TMEM132C)*, and cg11563844 (annotated to *STARD13* and *KL)*. In contrast, the other five CpGs (annotated to genes *SPG7, ANK1, MIR486, MYO1C, ABR, ALDH16A1*, and *FLT3LG*) showed independent evidence of association in our dataset (*p*-values ranging from 0.05 to 7.38E-04; Supplementary Table 6) with consistent effect directions and can therefore be regarded as a first independent replication of the results of Smith et al. (17).

### Half of the DNAm EWAS signals correlate with gene expression

To elucidate the potential functional implications of the DNAm associations highlighted above, we performed correlation analyses between the DNAm levels and corresponding mRNA expression data generated in the same individuals from the same tissue slices. To this end, we chose all 12 significantly associated CpGs from the EWAS meta-analysis results (i.e. both DMPs [Table 3, Table 4] and DMRs [Table 2]), as well as the eight available AD-associated CpGs in EC from Smith et al. (17) and correlated the DNAm levels with gene expression levels of the annotated gene(s) according to the Illumina manifest (v1.0 B5) for the EPIC array and the GREAT annotation tool. Within DMR “windows”, we selected the CpG showing the strongest association with AD case-control status for correlation with mRNA levels. Overall, this led to Spearman rank correlations of 39 DNAm-mRNA pairs (Supplementary Table 7). Ten of these pairs (with eight unique CpGs, two of which were CpGs from the Smith et al. EC meta-analysis (17)) showed evidence for a statistically significant DNAm-mRNA correlation after multiple testing correction (Table 5) accounting for 39 individual genes (Supplementary Table 8 and 9). In addition, using the genes with significant correlations as outcome in differential gene expression analyses performed on the same RNAseq data revealed that all ten were also significantly differentially expressed with respect to both AD case-control status and Braak stage (Table 5). One additional CpG-mRNA pair, which only showed a borderline negative correlation here (cg05972352 vs. ENSG00000126217; rho = -0.15, P-value = 0.061) was reported to show a significant correlation in the same direction in the study by de Witte et al, 2021 (40).

**Table 5:** Experiment-wide significant Spearman rank correlations between CpG DNAm and gene expression levels. Experiment-wide significant Spearman rank correlations between CpG DNAm and gene expression levels. β: Effect sizes of gene expression association with AD case-control status / Braak stage; P: P-value of gene expression association with AD case-control status / Braak stage after multiple testing adjustment with the Benjamini Hochberg method; Location: Location of the CpG in the genome with respect to the correlated gene; Results of all 39 tested DNAm-mRNA pairs can be found in Supplementary Tables 6, 7, and 8. One additional CpG-mRNA pair was found to significantly correlate in human brain samples by de Witte et al, 2021 (40); in our data, this pair showed a borderline significant correlation in the same direction as described by ref. (40) (see Supplementary Table 7).

| CpG | Gene | Spearman's $\rho$ | $P_{\text{Spearman}}$ | $\beta_{\text{Case / Control}}$ | $P_{\text{Case / Control}}$ | $\beta_{\text{Braak}}$ | $P_{\text{Braak}}$ | Location of CpG |
| --- | --- | --- | --- | --- | --- | --- | --- | --- |
| cg22090150 | CYB5D2 | -0.35 | 3.9E-04 | -0.28 | 0.000609 | -0.56 | 0.000512 | Intergenic |
| cg04520340 | PRKCZ | -0.28 | 5.59E-03 | -0.49 | 2.83E-11 | -0.98 | 2.83E-11 | gene body |
| cg22388948 | TENT5A | -0.28 | 5.59E-03 | 0.42 | 1.67E-07 | 0.79 | 6.69E-07 | gene body |
| cg05417607 | MYO1C | 0.26 | 1.17E-02 | 0.49 | 2.93E-12 | 0.93 | 2.83E-11 | gene body |
| cg22090150 | ANKFY1 | 0.24 | 1.76E-02 | 0.46 | 4.68E-09 | 0.88 | 2.03E-08 | gene body |
| cg05228284 | DIRAS1 | 0.24 | 1.76E-02 | -0.51 | 7.54E-12 | -1.10 | 5.85E-13 | 5'-UTR CGI |
| cg05066959 | ANK1 | -0.23 | 2.28E-02 | -0.39 | 6.07E-07 | -0.76 | 1.30E-06 | gene body island |
| cg20648333 | PALD1 | 0.22 | 2.97E-02 | 0.17 | 0.025071 | 0.35 | 0.015889 | gene body |
| cg20648333 | PRF1 | 0.22 | 3.03E-02 | 0.36 | 3.90E-06 | 0.60 | 0.000114 | Intergenic |
| cg20618448 | FLT3LG | 0.20 | 4.29E-02 | 0.36 | 1.67E-07 | 0.75 | 3.64E-08 | Intergenic |

Generally, the correlation coefficients only indicated moderate (maximal rho = -0.35), but statistically significant correlations between DNAm and mRNA expression in this dataset. The comparatively moderate extent of the correlations likely reflects the fact that gene expression is regulated by a number of other (epi-)genetic mechanisms beyond DNAm (37). Another noteworthy observation is that the signs of correlation coefficients of significant DNAm-mRNA pairs were both positive and negative, suggesting a complex relationship between DNAm status at these positions and their effect on mRNA expression. This is likely due to the fact that the majority of CpGs (nine out of ten) among the significantly correlated DNAm-mRNA pairs was located in gene bodies or distal to the stop-codon, while only one (CpG cg05228284) was located in an CpG-island (CGI) in the 5’ untranslated region (UTR). Classically, DNAm is considered a mark of transcriptional repression (here expected to elicit a correlation with a negative sign), however, this only applies to CpGs located in promoter CGIs, not necessarily those located elsewhere (38). Therefore, the presence of both positive and negative correlations between DNAm and mRNA levels (Table 5) is not unexpected.

### Epigenetic age acceleration is associated with AD in EC

In agreement with prior evidence, both DNAm age estimators were highly correlated with chronological age (HMTP: Pearson’s r = 0.56, *p* = 1.078E-13; CorCl: r=0.81, *p* < 2.20E-16) in our dataset. However, CorCl showed a (much) stronger correlation with chronological age compared to HMTP, and also did not show the tendency to under-estimate epigenetic ages compared to chronological age (Figure 3).

**Figure 3:**
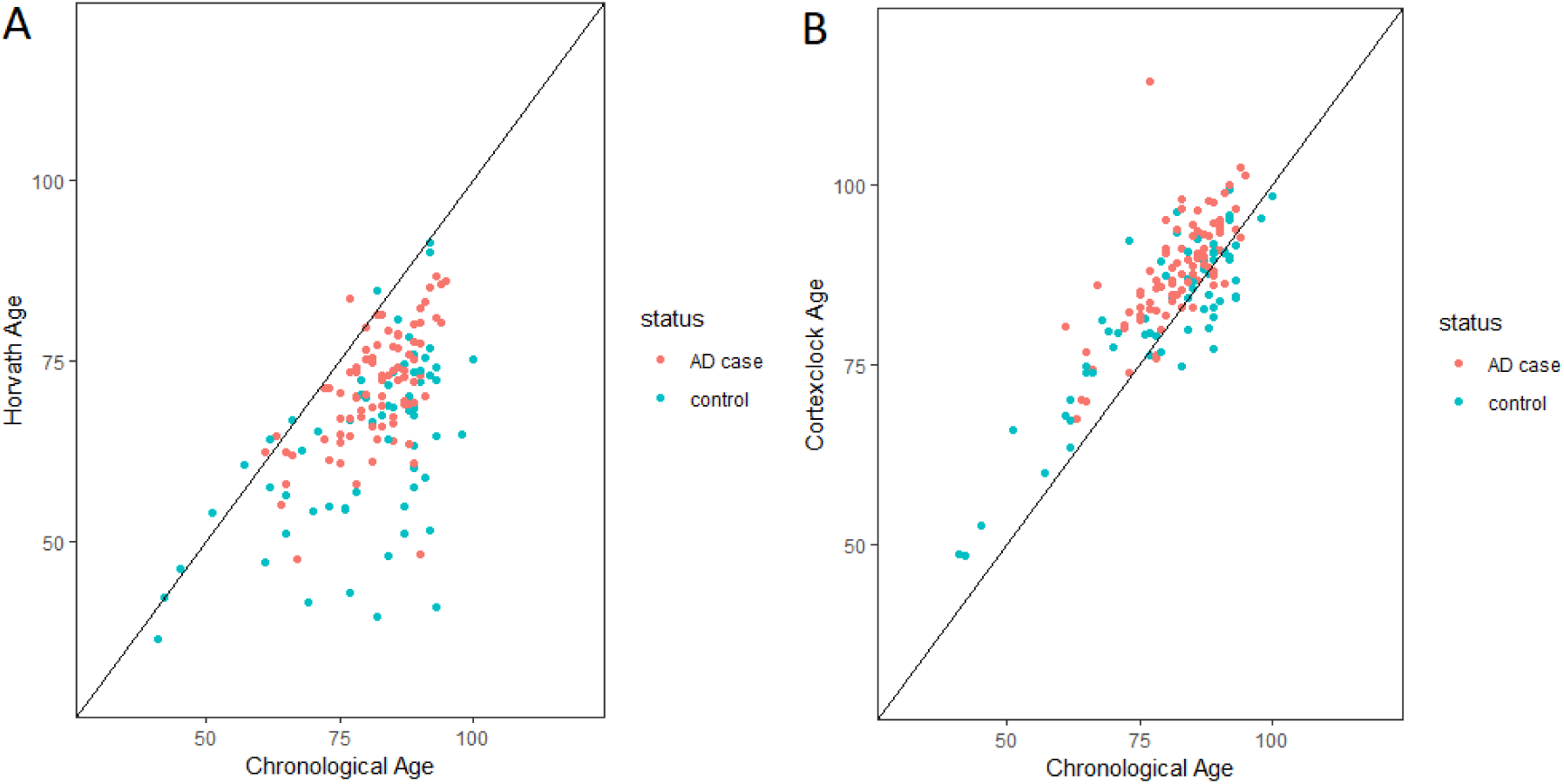
Scatter plots depicting the relation between chronological and epigenetic age colored according to AD case-control status. Panel A. Horvath multi-tissue age predictor (left); panel B. cortex clock (right); red: AD case; blue: control.

Using these estimates, we determined the degree of “age acceleration” which was defined as the residual from a linear regression of DNAm age on chronological age (39). This estimate was probed for an association with either AD case-control status or Braak stage using linear regression. Our expectation was that samples with an advanced disease state (e.g., AD vs. control, or high Braak stage vs. low Braak stage) would also show a more pronounced age acceleration (i.e., older epigenetic age when compared to chronological age). In concordance with this expectation, we found that age acceleration estimates were, indeed, associated with disease state (2.72E-05 ≤ *p* ≤ 1.96E-04, Table 6), with higher age acceleration being associated with AD cases or higher Braak stages (Table 6). In an additional model, we ran regression models similar to those for the primary EWAS, i.e. accounting for genetic ancestry as well as unknown confounders with respect to the DNAm data. After including these variables, only the CorCl remained significantly associated with AD case-control status (effect = 0.28, *p* = 4.70E-03), while some of the other estimates still showed suggestive evidence of association of age acceleration with disease state (Supplementary Table 10). Of note, DNAm PC1 and PC2 showed consistent associations with disease state, and therefore likely capture variance in DNAm data that reflect unknown confounders in the dataset (Supplementary Table 10), with the true level of association likely located somewhere between both models.

**Table 6:**
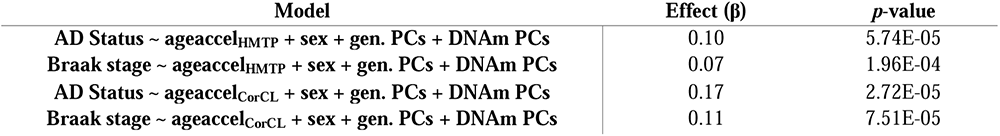
Results of linear regression analyses testing the association between epigenetic age acceleration and AD using the HMTP and CorCL DNAm age estimators. β: Effect sizes of AD case-control status / Braak stage with HMTP / CorCL age acceleration

| Model | Effect ($\beta$) | *p*-value |
| --- | --- | --- |
| **AD Status ~ ageaccel<sub>HMTP</sub> + sex + gen. PCs + DNAm PCs** | 0.10 | 5.74E-05 |
| **Braak stage ~ ageaccel<sub>HMTP</sub> + sex + gen. PCs + DNAm PCs** | 0.07 | 1.96E-04 |
| **AD Status ~ ageaccel<sub>CorCL</sub> + sex + gen. PCs + DNAm PCs** | 0.17 | 2.72E-05 |
| **Braak stage ~ ageaccel<sub>CorCL</sub> + sex + gen. PCs + DNAm PCs** | 0.11 | 7.51E-05 |

## Discussion

In this study we performed various EWAS analyses on a large collection of DNAm profiles generated in human EC tissue samples. Meta-analysis of newly generated DNAm data with those from two previous EC-based EWAS provides evidence for four novel loci showing significant association with either AD case-control status or Braak stage after stringent multiple testing correction. Using RNAseq data generated from the same individuals/tissue samples, we identified significant correlations between DNAm levels and mRNA expression for 10 out of the 39 DNAm-mRNA pairs within these EWAS loci. One additional locus (*MCF2L* [ENSG00000126217]) which only showed borderline significance here was recently reported to correlate with DNAm at cg05972352 in the study by de Witte et al. (40) and can arguably be counted as one additional relevant DNAm-mRNA pair in the context of our analyses. The most notable of our novel associations was observed with a CpG-probe (CpG cg22388948) in *TENT5A* (a.k.a. *FAM46A*), which not only showed consistent effect directions across all three analysed datasets, but also exhibited a significant (negative) correlation with mRNA levels of the same gene. *TENT5A* represents a promising novel AD candidate gene due to its previous association with several AD-relevant phenotypes by GWAS. Functionally, it belongs to the nucleotidyltransferase (NTase) fold superfamily (FAM46), which serve as non-canonical poly(A) polymerases involved in the modification of cytosolic and/or nuclear RNA 3’ ends and, hence, in the regulation of gene expression (41). Regarding the other novel signals of our EWAS, we note that although no direct evidence exists linking either *LIPE* (encoding lipase E [a.k.a. hormone sensitive lipase {*HSL*}], near CpG cg03073402 on chromosome 19p) or *PRF1* (encoding perforin 1; near CpG cg20648333 on chromosome 10q) to AD, both genes are involved in molecular pathways, i.e. lipid metabolism (42) and the immune system response (43), respectively, which are both highly relevant in AD pathogenesis based on recent GWAS data (44,45). Furthermore, the EWAS catalog lists both CpGs are associated with “human aging” (34), which further emphasizes the potential relevance of these loci. The possible link to AD pathogenesis of the last novel CpG-site cg05228284, near *DIRAS1*, is less obvious. This gene is highly expressed in the cerebellum, cortex, and frontal cortex according to GTEx. It belongs to the Ras superfamily of monomeric GTPases and has been previously reported as a tumor suppressor and is annotated to gene ontology pathways such as GTPase activity, protein binding, and signal transduction (46). Other relevant outcomes of our study are the independent confirmation of some, albeit not all, previous EC-based EWAS signals (17), and the observation that epigenetic age in EC is accelerated with increasing AD progression using two recently proposed estimators of DNAm age, a finding that is consistent with previously published data (47–49). We also confirm previous results that the HMTP (a.k.a. “Horvath clock”), which was trained on several tissues, may not be ideal to estimate DNAm age in the human brain cortex, and that, instead, the recently proposed “cortex clock” may be better suited for DNAm analyses in this tissue (30,50). One potential concern with these latter analyses is that the HMTP was originally derived using DNAm data from the 450K array (as opposed to the EPIC array used here). However, we note that the overlap between the CpG-probes present on the 450K and EPIC array is large, and it has previously been shown that the HMTP method provides reliable results when comparing data from both arrays (51).

The strengths of our study are its comparatively large sample size (n = 149 novel EC samples; n = 337 in the EWAS meta-analyses), the analysis of a brain region highly relevant for AD research (i.e. EC), the use of the hitherto highest resolution DNAm profiling microarray (i.e. the Methylation EPIC array featuring 850K CpG-probes), and the parallel availability of RNAseq-based mRNA expression data (allowing for detailed DNAm-mRNA correlation analyses). Despite these strengths, our study is also subject to a number of limitations which include the following. First and foremost, we used “bulk tissue” for DNAm profiling and RNAseq. Bulk tissue samples represent an agglomerate of different cell-types whose proportions (and DNAm and mRNA profiles) may vary across different samples (e.g. they may change as the disease progresses), a situation that may have affected the outcomes of our study. Single nucleus-based DNAm profiling and RNAseq would be required to fully address this question, which was not feasible in the context of our study. The next-best approach is to estimate sample-specific cell-type proportions using cellular deconvolution (ideally based on reference data from the same tissue with known cell-type composition) and include these estimates into the analyses. To this end, we estimated cell-type composition based on DNAm reference data obtained from human prefrontal cortex samples (to the best of our knowledge, no DNAm reference data exist for EC) using the *estimateCellCounts* function in the R package Minfi (52) and regressed these out with the *removeBatchEffect* function in the R package limma (53). Using this approach did not appreciably change our results and none of our top signals changed in the AD case-control or Braak stage EWAS (Supplementary Tables 1 and 2). Second, our reanalysis of publicly available DNAm profiles from EC resulted in slightly increased (i.e. less significant) P-values when compared to those reported in the primary publications (17). This is likely due to a more conservative covariate adjustment scheme used in our data processing workflow. Repeating all meta-analyses based on a covariate adjustment paradigm similar to that used in Smith et al. (17) (i.e. include DNAm covariates until λ drops to <1.2 in each individual dataset) led to an overall increase in statistical significance of most of the meta-analysis results (Supplementary Table 11), approaching those previously reported. We further note that Smith et al. (17) did not adjust their results for genetic ancestry, which may have also affected their test statistics. Therefore, the “true” level of statistical support likely lies somewhere in between both approaches, and suggests that the EWAS results reported here are likely conservative. Third, our meta-analysis is based on fewer CpGs (n = 304,996) as reported by the overlap of the London-1 and London-2 datasets (up to n = 403,763 (7,11)). This is due to the fact that we used a different DNAm microarray (“MethylationEPIC”) here, which does not show perfect overlap in CpG-probes with the predecessor array (“450K”) and removed a comparatively large number of probes from all datasets during QC. Fourth, as previously noted (17), EWAS meta-analysis tend to show inflations of the test statistics, which we also observed here. One possible reason is heterogeneity of the effect estimates across studies due to technical reasons. To address this issue, we repeated the meta-analysis using random-effect models, which indeed showed a much less pronounced degree of inflation than fixed-effect models (Case-control: λ_fixed-effect_ = 1.16, λ_random-effect_ = 0.92; Braak stage: λ_fixed-effect_ = 1.24, λ_random-effect_ = 1.00; Supplementary Figure 5). However, we note that for only two out of the twelve experiment-wide significant DMPs, heterogeneity measured by the i^2^ statistic exceeded 50% (Supplementary Table 12), supporting the appropriateness of using the fixed-effect models, as was done in other recent EWAS meta-analyses (17). Fifth, we note that our samples displayed relatively low bisulphite conversion efficiency rates necessitating to lower the exclusion threshold from 80%, which is typically used, to 65% here. To ensure that the EWAS results obtained were not driven by artifacts due to low sample quality, we repeated our meta-analyses using 80% as threshold, which did not substantially change the results (Supplementary Table 13). While several CpG-sites now showed slightly larger p-values, these can most likely be attributed to a loss in power due to a smaller sample size (300 [i.e. minus 37] and 280 [i.e. minus 40] for the Braak stage and case-control analyses, respectively). Sixth, we are not able to discern cause-effect relationships from our data. The observed epigenetic associations may reflect molecular changes preceding (and perhaps modifying) the underlying disease process, but they may also represent a *consequence* of the same. However, the inability to disentangle the sequence of events is not a limitation of our EWAS, but of any epigenetic study. Notwithstanding, our novel results imply several new loci and potential molecular mechanisms that are associated with AD and, as a result, provide important new insights to our understanding of the pathogenetic mechanisms underlying the disease process. Lastly, despite going to great lengths to minimize spurious associations by applying conservative QC thresholds in all steps of our data processing steps, we cannot exclude the possibility that some undetected confounding factors have impacted our results. However, the overall high concordance between our and previous EWAS results argues against a strong systematic bias specific to our data and/or results. Nevertheless, all novel results to emerge from this work should be considered preliminary until independent validation is reported.

In summary, in the largest AD EWAS performed on EC samples to date, we identified a total of 12 epigenome-wide significant CpGs, four of which are novel. Six of these CpGs show significant correlations with corresponding mRNA levels in the same samples, highlighting their potential downstream effects on gene expression. Future work is needed to validate our findings and to clarify the role of the newly implied loci in AD pathogenesis.

## Acknowledgements

This work was supported by the Cure Alzheimer’s Fund (as part of the “CIRCUITS” consortium) to L.B., the EU Horizon 2020 Fund (as part of the “Lifebrain” consortium, #732592) to L.B., and by the Deutsche Forschungsgemeinschaft (DFG) and the National Science Foundation China (NSFC) as part of the Joint Sino-German research project (“MiRNet-AD”, #391523883) to L.B. Additional support was provided by the DFG Research Infrastructure NGS_CC (project 407495230) as part of the Next Generation Sequencing Competence Network (#423957469). NGS analyses were carried out at the Competence Centre for Genomic Analysis (Kiel). We acknowledge the Oxford Brain Bank, supported by the Medical Research Council (MRC), Brains for Dementia Research (BDR) (Alzheimer Society and Alzheimer Research UK), Autistica UK and the NIHR Oxford Biomedical Research Centre. Last but not least, we acknowledge the high-performance compute environment (“OmicsCluster”) at University of Lübeck where most data processing and analysis steps of this study were run.

## Author contributions

Design of the study, supervision, and acquisition of funding: LB. Ascertainment of brain tissue and neuropathological examinations: LP. Handling of tissue samples and generation of molecular data: YS, VD, TW, SSS, AF, JF, SF. Data processing and statistical analyses: YS, MS, OO, CML. First draft of the manuscript: YS, LB. Critical revision and final version of manuscript: all authors.

