## Supplementary Material for "Entorhinal cortex epigenome-wide association study highlights four novel loci showing differential methylation in Alzheimer’s disease"

**Running title:** Alzheimer’s disease EWAS in entorhinal cortex

**Keywords:** Alzheimer’s disease; DNAm; brain; entorhinal cortex; meta-analysis; methylation EWAS

### Supplementary Methods

#### DNA and RNA extraction and processing

For 183 entorhinal cortex (EC) samples, genomic DNA and total RNA were extracted from approximately 50mg and 25mg of frozen tissue, respectively. DNA extraction was performed using the DNeasy Blood & Tissue Kit (Qiagen, Hilden, Germany), while for total RNA, we used the mirVana miRNA kit (Thermo Fisher Scientific, USA). All extraction steps were carried out according to manufacturer’s instructions. DNA and RNA were quantified using a NanoDrop ONE spectrophotometer (Thermo Fisher Scientific). In addition, RNA integrity was assessed using a Bioanalyzer 2100 instrument and the RNA 6000 Nano LabChip kit (Agilent Technologies, USA).

#### EPIC array profiling

DNA methylation (DNAm) profiling was performed using the “Infinium MethylationEPIC” array (Illumina, Inc.) on aliquots of DNA extracts diluted to ~50ng/µl concentration. The EPIC array has been extensively validated and obtained DNAm values were highly correlated with values obtained by other methods, such as the 450K array, methylation capture sequencing, and whole-genome bisulphite sequencing (1–3). DNA samples were subjected to bisulphite conversions using the EZ DNA Methylation kit (Zymo Reasearch) following the alternative incubation conditions for the Illumina Infinium MethylationEPIC Array from the supplier, and then hybridized to the EPIC array and scanned on an iScan instrument (Illumina, Inc.) according to the manufacturer’s instructions (Document#1000000077299v0). All DNA samples were treated in consecutive laboratory experiments to minimize the potential for batch effects. Raw DNAm intensities were called using the iScan control software (v2.3.0.0; Illumina, Inc.) and exported in .idat format for downstream processing and analysis.

#### DNA methylation data processing and quality control

DNAm data pre-processing and quality control (QC) was performed in R (v. 3.6.1) using the package bigmelon with default settings (4), unless otherwise noted. Idat files were loaded into R and β-values were calculated according to the following formula, with *I*_met_ being the intensity of the methylated signal, and *I*_ume_ being the intensity of the unmethylated signal.

(1) β = *I*_met_ / (*I*_met_ + *I*_ume_ + 100)

Samples were excluded from the analysis if (a) the bisulphite conversion efficiency of the sample according to the *bscon* function in the bigmelon package was below 65%, (b) the sample had a beadcount < 3 in more than 5% of all probes, (c) the sample had a detection p-value below 0.05 in more than 1% of all probes, (d) the sample was identified as an outlier according to the *outlyx* function in the bigmelon package using a threshold of 0.01, (e) the sample showed a large change in β-values after normalization according to the *qual* function in the bigmelon package with a threshold of 0.1, (f) the sample showed a discrepancy between predicted sex according to the Horvath multi-tissue epigenetic age predictor (5) and reported sex, or (g) there was a greater than 70% discrepancy between genotypes of 42 SNPs determined concurrently from the EPIC and GSA SNP genotyping array (see below). All samples were normalized with the *dasen* function of bigmelon.

Cytosine-phosphate-guanine (CpG) probes were removed from the analysis if (a) they had a detection *p*-value below 0.05, (b) they had a beadcount < 3, (c) they aligned to multiple locations according to Nordlund et al. (6), (d) they aligned to locations influenced by SNPs according to Zhou et al. (7), or (e) they aligned to locations on the X and Y chromosomes.

The final analyses included 665,796 CpG-probes in 149 samples for the Alzheimer’s disease (AD) case-control epigenome-wide association study (EWAS) and 142 samples for the Braak stage EWAS. A detailed sample description can be found in Table 1.

#### RNA sequencing and quality control

Total RNA was extracted from EC samples as described above. Libraries were generated using the TruSeq Stranded Total RNA kit (Illumina) and sequenced at a target depth of 30 M reads per samples in 2x100 bp paired-end mode on a NovaSeq 6000 device (Illumina) with 2 x 100 bp reads. All transcripts were pseudoaligned to the human transcriptome using the Ensembl human genes annotation (version 100) (8) and were quantified using kallisto (9) and the –bias and –rf-stranded options. The resulting trancriptome profiles were filtered for protein coding and lncRNA genes only and summarized to the gene level (summing up read counts of all isoforms of each gene) and normalized using DESeq2 (10).

#### Batch effect correction for DNAm data

We performed a principal component analysis (PCA; using the R base function *prcomp*) on a subset of uncorrelated CpGs to capture variation in DNAm irrespective of phenotype. To this end, we created a set of uncorrelated CpGs by dividing the genome into 100kb bins and followed by random selection of one CpG-probe per bin. In the full model, 13 such “DNAm PCs” were included as covariates in the EWAS analyses.

To account for differences in genetic ancestry we performed a PCA (using PLINK v1.9 “--pca”) on an LD-pruned set of SNP markers (--indep-pairwise 1500 150 0.2) derived from genome-wide SNP genotyping data generated in parallel on the same DNA samples of each individual using the Global Screening Array (GSA; Illumina, Inc). The first 20 genetic ancestry PCs were used as covariates in the EWAS analyses. More details on the genotyping and QC procedures can be found in a previous publication (11).

#### Alzheimer’s disease poly-epigenetic scores

To assess the correspondence of our novel DNAm data to previous EWAS on the topic, poly-epigenetic scores (PES) for each individual were calculated based on the test statistics from two publicly available AD EC datasets (GEO accession numbers GSE59685; GSE105109). To this end, we combined uncorrelated CpGs into one aggregated DNAm variable and tested these as predictors in regression models analogous to the primary EWAS. The descriptions of the datasets used for PES calculations can be found in the primary publications (12,13). For the PES calculations, test statistics from the meta-analysis with varying *p*-value thresholds (*p* < 1, *p* < 1.00E-04, *p* < 1.00E-05, *p* < 1.00E-06, *p* < 1.36E-07) were used. For each *p*-value threshold, we created a set of uncorrelated CpGs by dividing the genome into 100kb bins and followed by the selection of the CpG with the most extreme effect size per bin. Here, the effect sizes from the meta-analysis for London-1 and London-2 (*β*) and the normalized DNAm values (*CpG*) for each of the *n* uncorrelated CpGs for each *p*-value threshold were used and combined as follows:

$$\sum_{i=1}^{n} \beta_{i}{*CpG}_{i}$$

PES-based association analyses used linear regression models to predict Braak stage as outcome and PES as predictor, adjusting for the same covariates as in the primary EWAS analyses.

#### Epigenetic age estimation

Two epigenetic age predictors were used in our analyses: 1. the “Horvath multi-tissue predictor” (HMTP) (5) and 2. the “cortex clock” (CorCl) (14). Since most other popular epigenetic clocks (Hannum (15), PhenoAge (16), GrimAge (17)) were calibrated for blood tissues, we did not include analyses of these age estimators in this study. DNAm age using the “Horvath multi-tissue predictor” HMTP was calculated with the R script provided in Horvath (5), using the DNAm raw data (with prior removal of probes failing QC). Prior to the age estimation, this algorithm uses BMIQ as data normalization method. For the “cortex clock” (CorCl), the normalized data from the processing pipeline described above were used directly in code provided by the authors of ref. (14) (<https://github.com/gemmashireby/CorticalClock>). Age acceleration was defined as residual from a linear regression of epigenetic age on chronological age.

#### DNA-mRNA correlation analyses and differential gene expression analyses

The normalized RNA-seq data for the selected CpG candidate genes were correlated to their corresponding DNAm signal, using the Spearman method in R’s *cor.test* function. The resulting *p*-values were corrected for multiple testing using the Benjamini-Hochberg procedure (as implemented in R’s *p.adjust* function). Prior to the correlation analyses, outliers (>1.5 * inter-quartile-range below/above 1st/3rd quartile) were removed. Additionally, linear models were fit (using R’s *glm* function with family = “gaussian”) to predict the (log-transformed) gene expression estimates from the corresponding DNAm values. The following potential confounders were included as co-variables in these models: age at death, sex, RNA integrity (RIN) and post-mortem interval (PMI). All continuous variables were scaled and centred before fitting the model. Finally, additional models were fit adding disease status (AD case vs. control) or Braak stage (as a continuous variable), respectively. The corresponding F-statistic was used to assess the significance of the (additional) amount of gene expression variance by the predictors of interest. As for the correlation analysis above, multiple-testing was addressed by false-discovery-rate (FDR) estimation as per Benjamini-Hochberg.

### Supplementary Figures

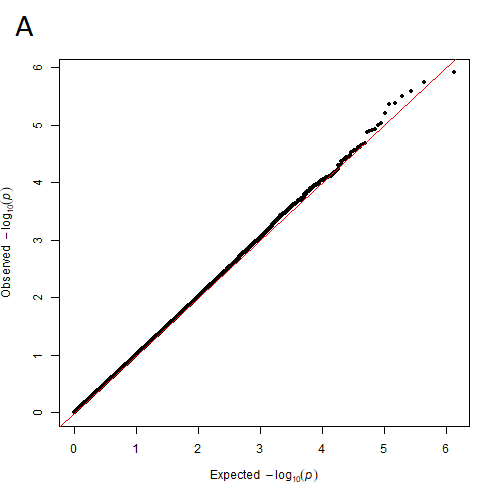

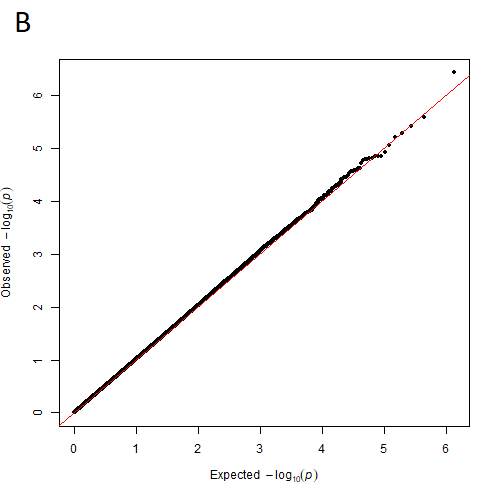

S1: QQ-plots for the results of the AD case-control EWAS (A, λ=1.0005), and the Braak stage EWAS (B, λ=1.0175) in the Oxford datasets.

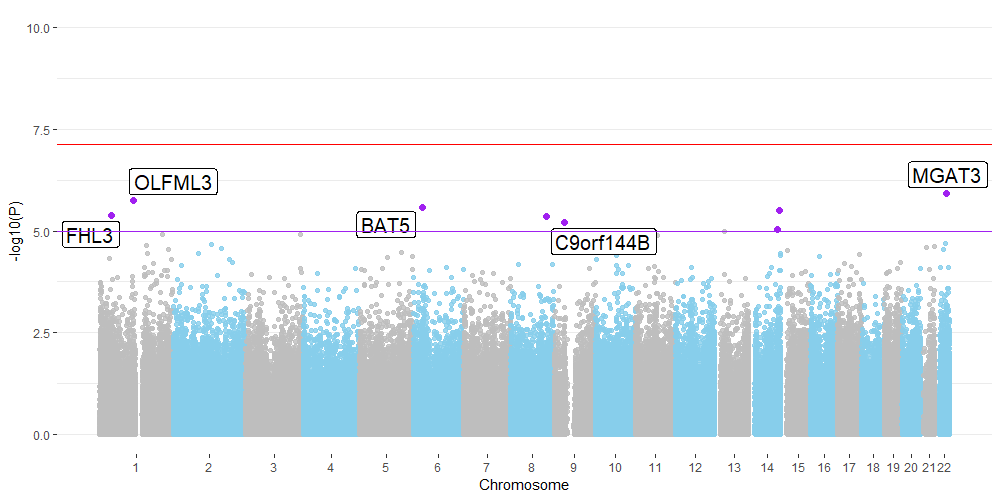

S2: Manhattan plot for the results of the AD case-control EWAS in the Oxford dataset; the red line indicates the experiment-wide significance threshold of 7.51E-08, whereas the purple line indicates the suggestive significance threshold of 1.00E-05. CpGs with suggestively significant association are marked in purple and annotated with the gene name according to the Illumina manifest (v1.0 B5). NB that three CpGs (on chr. 8 and chr. 14 were not annotated to any genes).

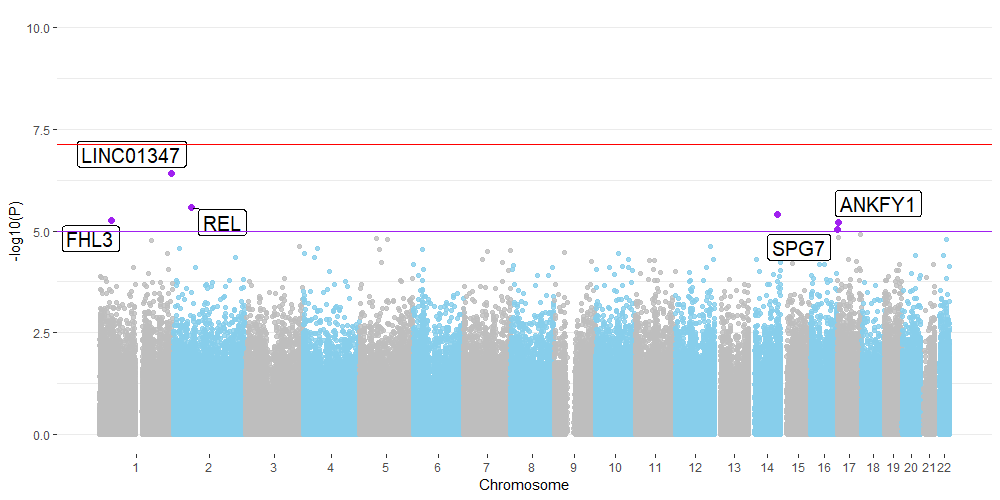

S3: Manhattan plot for the results of the AD Braak stage EWAS in the Oxford dataset; the red line indicates the experiment-wide significance threshold of 7.51E-8, whereas the purple line indicates the suggestive significance threshold of 1.00E-05. CpGs with suggestively significant association are marked in purple and annotated with the gene name according to the Illumina manifest (v1.0 B5). NB that one CpG (on chr. 14 was not annotated to any genes).

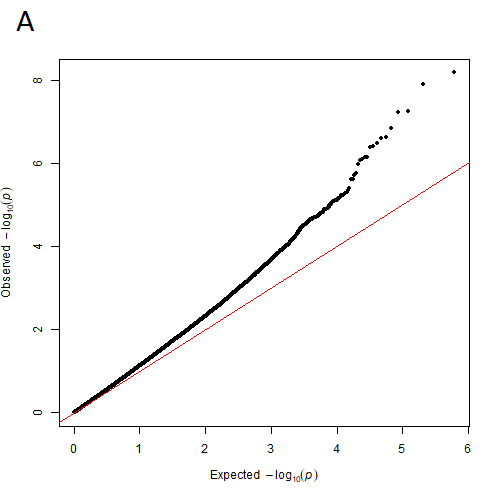

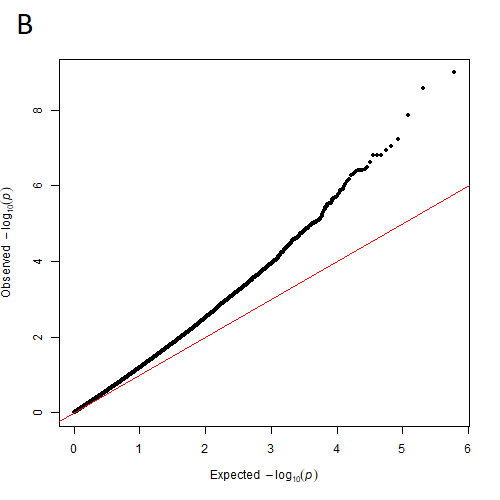

S4: QQ-plots for the results of the AD case-control meta-EWAS (A, λ=1.16), and the Braak stage meta-EWAS (B, λ =1.24) with fixed-effect models across three datasets (London-1, London-2, Oxford).

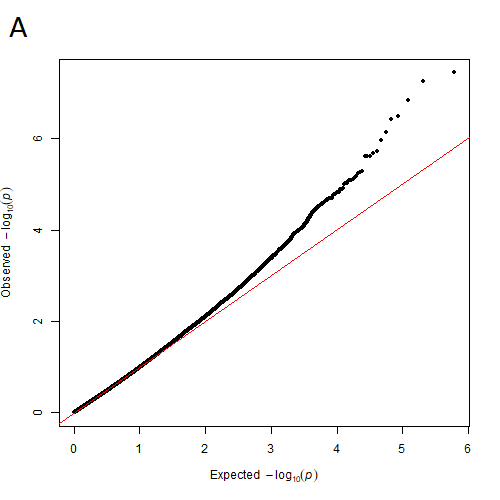

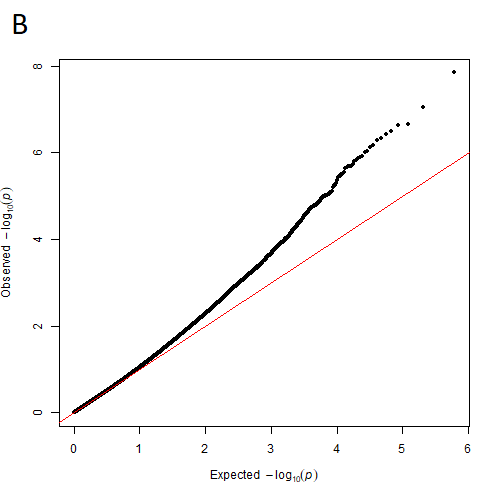

S5: QQ-plots for the results of the AD case-control meta-EWAS (A, λ=0.92), and the Braak stage meta-EWAS (B, λ =1.00) with random-effect models across three datasets (London-1, London-2, Oxford).
